# YAEL: Your Advanced Electrode Localizer

**DOI:** 10.1101/2023.08.04.552023

**Authors:** Zhengjia Wang, John Magnotti, Xiang Zhang, Michael S. Beauchamp

## Abstract

Intracranial electroencephalography (iEEG) provides a unique opportunity to measure human brain function with implanted electrodes. A key step in neuroscience inference from iEEG is localizing the electrodes relative to individual subject anatomy and identified regions in brain atlases. While there are number of workflows for electrode localization, most suffer from one or more limitations. The first limitation is a lack of integration: scientists must install and use different software packages for each localization step. Second, they are inefficient: while most iEEG analysis steps can be automated, electrode localization is still largely a manual process. Third, most current tools are limited to the localization process itself, leaving users without the ability to create high-quality visualizations for clinical and research purposes. We developed YAEL (Your Advanced Electrode Localizer) to overcome these limitations. First, YAEL is completely integrated: a single easy-to-use graphical user interface (GUI) controls every step of the localization process. Second, YAEL uses a flexible 3D viewer and automation tools to make accurate localization of electrodes quick and easy. Third, after localization is complete, YAEL leverages the same viewer to create high-quality visualizations of electrode data including identified brain areas from atlases; the response to experimental tasks measured with iEEG; and clinical measures such as epileptiform activity or the results of electrical stimulation mapping. YAEL contains more than 30,000 lines of code, is free and open source, and can be installed in minutes on Mac, Windows and Linux platforms from *https://yael.wiki*. User interactions with YAEL occur through a web browser ensuring a familiar user experience and consistent operation across platforms and whether YAEL is used locally or deployed in the cloud.

## Introduction

Intracranial electroencephalography (iEEG) is a powerful technique in human neuroscience that records the activity from small groups of neurons from implanted electrodes implanted in the brain. A critical step in the analysis of iEEG data is defining the anatomical location of each electrode accurately and efficiently. Accurate electrode localization is critical for neuroscience inference, but efficiency is also important, as a single patient may be implanted with hundreds of electrodes.

The essential steps in electrode localization are straightforward. Typically, structural magnetic resonance imaging (MRI) scans are collected before the implantation surgery. After surgery, a CT (computed tomography) scan is obtained. The post-operative CT and pre-operative MRI are then aligned. Electrode locations are identified using the CT (metal electrodes are easily localizable as high-intensity regions in the CT, but produce dark susceptibility artifacts in the MRI), then visualized on the MRI due to its superior anatomical contrast. Because of its importance, there are a plethora of tools for iEEG electrode localization, including iELVis (Groppe et al., 2017), ALICE (Branco et al., 2018), img_pipe (Hamilton et al., 2017), LeGUI (Davis et al., 2021), iEEGview (Li et al., 2019), BFM Tool (Wang et al., 2016), DELLO (Zhao et al., 2023), iElectrodes (Blenkmann et al., 2017), IELU (LaPlante et al., 2017), EpiTools (Medina Villalon et al., 2018), and iEEG-recon (Lucas et al., 2023). There are also tools designed for localizing deep brain stimulation (DBS) electrodes, such as Lead-DBS (Horn et al., 2019; Horn and Kühn, 2015), although compared with iEEG, DBS patients are implanted with many fewer electrodes that are exclusively subcortical and are only for stimulation rather than recording.

Given the numerous existing tools, what is the impetus for yet another electrode localizer? A major limitation of some existing methods is that they require scientists to install and learn several different software tools, written in different languages at different time by different groups using different data formats. Some packages, such as CURRY, are entirely commercial, charging thousands of dollars per license. Other software is freely available but relies on the commercial Matlab package and add-on toolboxes purchased at additional expense. For example, before using iELVis, users must purchase a license and install Matlab, then separately download and install the Matlab routines in the legacy (unsupported) version of BioImage Suite (Papademetris et al., 2006), the Matlab iELVis codebase, and MRIcroGL (Rorden and Brett, 2000). To actually localize electrodes, a complex series of steps in each different tools must be undertaken (click [here](https://openwetware.org/wiki/Beauchamp:iELVIS) for sample workflow). YAEL streamlines the process so that users can localize electrodes within a single GUI that is free from reliance on commercial software.

A second limitation of some existing methods is that they are inefficient. Recently, clinical iEEG practice has transitioned from subdural electrodes, in which grids or strips are placed on the surface of the cortex, to stereotactic EEG (sEEG), in which many electrode shafts are inserted into the parenchyma. Although many older localization tools have features tailored to electrode grids, YAEL fully supports both subdural and sEEG electrodes. YAEL’s flexible 3D HTML WebGL-based brain viewer gives full control over dozens of visualization parameters. This control is especially important for sEEG, as it can be difficult to differentiate which electrodes belongs to which shafts. To speed the localization process, YAEL provides tools for both interpolation and extrapolation so that it is not necessary to manually select all electrodes on a shaft. Refinement tools ensure that manually selected electrode locations are located precisely at the center of the corresponding CT density. Tutorial videos (available [here](https://openwetware.org/wiki/YAEL:Help#YAEL_Videos)) help new users get up to speed and the software download includes a sample dataset with a pre-implant MRI and a post-implant CT.

A third limitation of existing methods is that they offer limited utility: electrode locations must be exported to another package for further analysis and visualization. While YAEL can also function as a stand-alone localizer, a more powerful alternative is to use the flexible viewer in YAEL to make publication-quality images and movies using exactly the same GUI as for electrode localization. This flips the script on the traditional workflow, in which electrode locations are *exported* from the electrode localizer to another program. Instead, one of the many existing packages for iEEG data analysis, such as EEGLab (Delorme and Makeig, 2004), MNE-Python (Rockhill et al., 2022), FieldTrip (Oostenveld et al., 2011), RAVE (Magnotti et al., 2020), or a laboratory’s own software pipeline, can be used to calculate values for each electrodes. Then, these values are *imported* into YAEL for high-quality visualization. Figure 1 shows the outcome of this process, a combination of electrode, brain, and processed iEEG data combined with YAEL; Figure 4 shows additional visualizations for a variety of research and clinical use cases.

**Figure 1.**
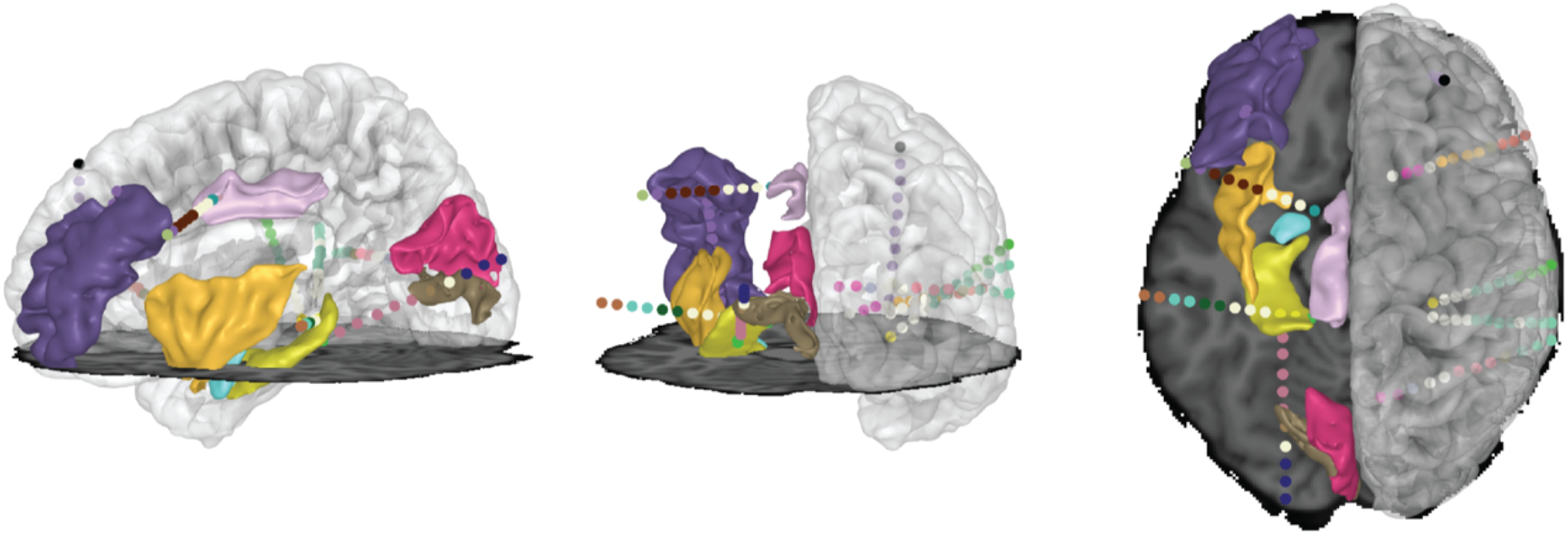
Highlights of the YAEL 3D viewer. Users can freely rotate the brain to show lateral view (left panel), posterior view (middle panel) or top view (right panel). Gray scale shows MRI data (horizontal plane shows axial slice through MRI data; right hemisphere shows translucent cortical surface model). Colored volumes show different anatomical regions of interest. Colored spheres show electrode contacts on different sEEG shafts. Color scale set by user and can reflect anatomical location and categorical or continuous experimental or clinical results (see Figure 4).

## Results

Figure 2 provides an overview of the YAEL workflow, divided into the major steps in the workflow: image inputs; pre-processing; electrode localization; and data visualization. YAEL is designed to function using the computational resources available in any scientific laboratory (a PC, Mac or Linux machine with any web browser). The heart of YAEL is the 3D brain viewer, programmed in HTML and JavaScript, with WebGL enabling hardware acceleration. The brain viewer is incorporated into a GUI written in R (R Core Team, 2023) with shinyR (Chang et al., 2023) extensions. All user interactions occur through a web browser, ensuring a consistent user experience across platforms.

**Figure 2.**
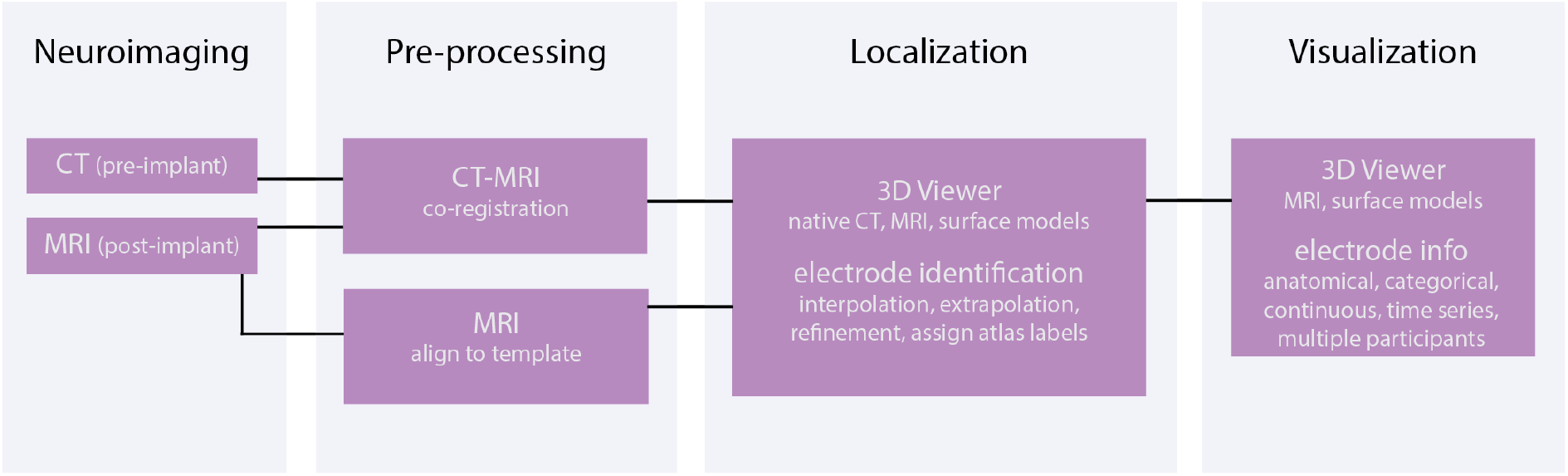
Flowchart of YAEL workflow.

### Inputs

YAEL provides a file chooser to make it easy for users to specify the location of the required MRI and CT datasets, accepted in both NIFTI and DICOM formats. The user then provides an electrode plan specifying the total number of electrodes and how they are grouped into different sEEG shafts and ECoG grids or strips.

### Pre-processing

Installing YAEL automatically installs two popular tools for co-registering the MRI and CT datasets, Advanced Normalization Tools (ANTs; Tustison et al., 2021) and NiftyReg (Modat et al., 2014). With a single click within YAEL, the datasets are aligned. For users that already have the extensive FMRIB software library (FSL) installed (Woolrich et al., 2009), YAEL also supports one-click registration with FLIRT (Greve and Fischl, 2009; Jenkinson et al., 2002).

Regardless of the registration tool selected, it is important to verify the CT-MRI alignment. YAEL facilitates this by providing tools to visualize the CT overlaid on the MRI dataset. Fine-tuning the co-registration is provided via modifiable command line parameters (specific to the tool being used), adjusted from within the YAEL GUI. If necessary, users can also use pre-aligned CT/MRI data, bypassing co-registration. However, because typical in-plane resolution is 0.3 mm for CT *vs*. 1 mm for MRI, using pre-aligned data that has been down-sampled in resolution may lead to poor localization results. For instance, smaller electrodes may be lost (Supplementary Figure 1). For this reason, YAEL uses the full-resolution CT during localization, rather than down-sampling it to the MRI resolution.

### Brain location identification

To identify the anatomical location of electrodes, YAEL supports aligning MRI data to a template brain. For instance, YAEL can use the included ANTs toolset to align the patient’s MRI data to the Montreal Neurological Institute template (Evans et al., 1993). Alternatively, for users that have the FreeSurfer neuroimaging toolkit installed (Dale et al., 1999; Fischl et al., 1999), YAEL can use the cortical surface model of the patient’s brain and align it to a surface-based template, allowing dozens of cortical areas to be automatically labeled (Desikan et al., 2006; Destrieux et al., 2010; Fischl et al., 2004). YAEL can call FreeSurfer with a single click and automatically load the resulting cortical surface models for improved visualization.

### Electrode localization

Identifying electrode locations is the most important and time-consuming aspect of the localization workflow, and YAEL provides several tools to improve the accuracy and efficiency of identifying electrodes. The most important tool is the sophisticated 3D viewer, which provides simultaneous visualization of 3D cortical surface models and 2D anatomical slices together with CT data and localized electrode positions (Figure 3A). Users can rotate the brain using the mouse or keyboard shortcuts, adjust the transparency of cortical and subcortical surface models, and display any combination of axial, sagittal and coronal views of the MRI dataset (MRI is displayed in grayscale with an adjustable solid color overlay for the CT). With a 2D viewer, it can be difficult to determine which electrodes should be assigned to which shaft. In contrast, with YAEL’s 3D Viewer, the spatial orientation of the electrodes in the same dataset is immediately apparent.

**Figure 3.**
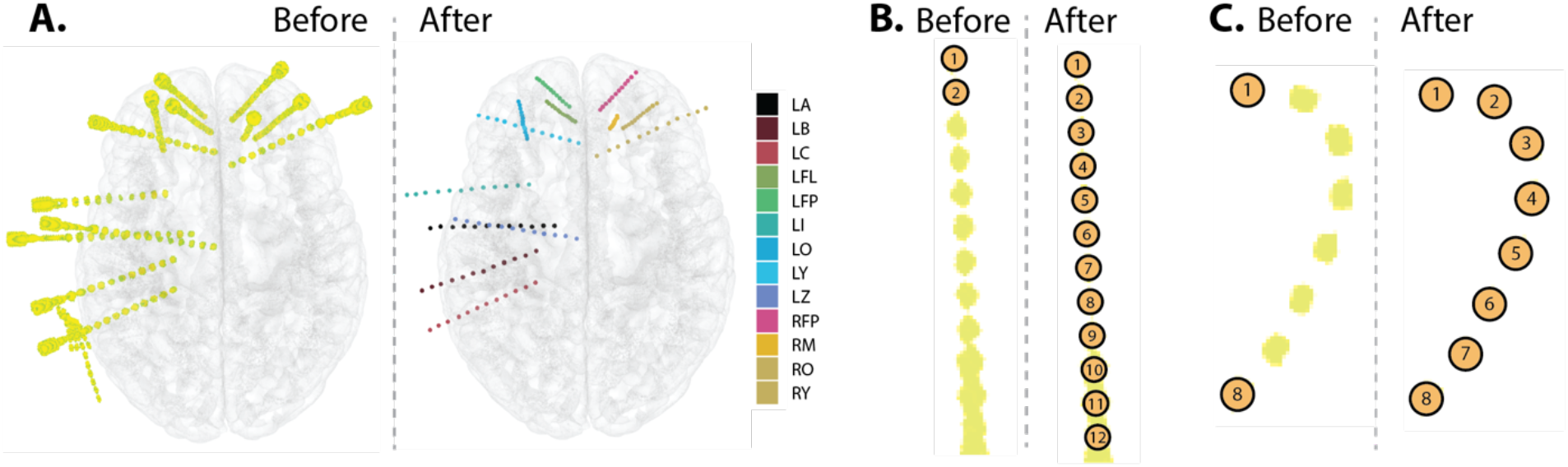
A. During localization (left panel), the viewer allows flexible 3D manipulation and visualization of the CT dataset (yellow color) and the MRI dataset (transparent cortical surface models). After localization (right panel), each electrode contact is visible as a sphere, with contacts on each sEEG shaft in a different color (the legend shows the code assigned to each shaft during implantation surgery). B. For automatic extrapolation, the user clicks the first two contacts of an sEEG shaft (shown) or a subdural electrode strip, then YAEL automatically localizes the remaining electrodes. C. For automatic interpolation, the user clicks on the first and last contact on a subdural electrode strip (shown) or an sEEG shaft. Clicking the “interpolate” button automatically localizes the intermediate electrodes. Interpolation succeeds despite the sharp curvature generated as the subdural strip conforms to the occipital pole.

Double clicking in the vicinity of a CT density in either the 3D viewer or an MRI slice view creates a new electrode, visible as a sphere, and applies an automatic refinement process to make sure that electrodes are positioned at the contact center by weighting the nearby CT densities. The location of the sphere can be manually adjusted with the mouse or keystroke for finer adjustment. The electrode plan provided by the user is used to populate a table with the locations and atlas labels of each contact. A fully manual mode is also available, allowing users to simply select locations on the cortical surface or the MRI volume in situations where a CT is not available, such as for intraoperative iEEG cases.

### Automation

YAEL expedites the process of electrode localization with two automation tools. The first automation tool is extrapolation (Figure 3B). The user selects the first two contacts in a shaft and clicks the “extrapolate” button—YAEL automatically assigns locations to the intermediate contacts, populating the electrode table. For interpolation (Figure 3C), the user selects two contacts at either end of an sEEG shaft, clicks the “interpolate” button, and then YAEL automatically identifies the intermediate electrodes. The number of extrapolated or interpolated electrodes can be automatically determined from the electrode plan or specified manually.

To account for any bends in the electrode shaft, YAEL’s interpolation feature iteratively performs interpolate and extrapolate operations. The distance between the two most recently created contacts is measured to determine the likely location of the next contact, but because electrode arrays are not precisely linear (due to shaft bending for sEEG electrodes and curvature of the cortical surface for ECoG electrodes) the precise location often differs from the computed location, so YAEL sequentially examines nearby (< 2mm) CT voxels to identify the precise position of the contact (CT voxels near previously selected contacts are excluded to avoid duplicate selection). This iterative approach is effective, even with subdural strips that have substantial bends around the occipital pole, as shown in Figure 3C.

### Interoperability

YAEL is designed to easily interoperate with other software with data import and export. Electrode data is stored in a simple plain text data table with one row per electrode and one column per field of data. The traditional workflow for electrode localization software is to *export* the co-ordinates (in MRI space or standard space) of each electrode for display in another package. YAEL supports this workflow: with a single click, a table with one row per electrode is saved to disk. The saved data for each electrode consists of the co-ordinates in native space and atlas space, and the atlas labels for each electrode.

An alternative mode for YAEL usage is to visualize the data using YAEL’s flexible 3D viewer. In this mode, the user *imports* data about each electrode into YAEL visualizes it using the 3D viewer. To make it easy for user to import data, YAEL automatically creates a template table with one row for each electrode. The user provides as many columns of data for each electrode as desired; YAEL intelligently uses the column label for visualization and ignores table cells with no data.

Electrode data can be generated manually by the user using common table editors or spreadsheet (such as Microsoft Excel) or generated algorithmically from another software package. In particular, there are many powerful analysis packages for analyzing the voltage by time data that constitutes iEEG, such as EEGLAB (Delorme and Makeig, 2004), FieldTrip (Oostenveld et al., 2011), EpiTools (Medina Villalon et al., 2018), MNE-Python (Rockhill et al., 2022), and RAVE (Magnotti et al., 2020). Many iEEG labs develop their own suite of semi-custom routines written in Python and Matlab for doing advanced analysis. YAEL can easily import data from any of these workflows for display.

### YAEL Visualization

The utility of importing data into YAEL (rather than exporting electrode locations to another program, the usual workflow for electrode localizers) is shown in five sample usage scenarios in Figure 4. In the first scenario, YAEL is used to visualize anatomical-functional information about each electrode available from brain atlases (Figure 4A). YAEL automatically labels electrodes by their anatomical identification. Clicking on an electrode displays all the anatomical-functional information available about an electrode, along with the atlas the information is derived from. By applying the atlases aligned to the individual subject MRI in the pre-processing stage (described above), users can select electrodes based on their anatomical identification. Users can select any combination of ROIs, and electrodes that do not fall in a desired ROI are colored gray. In this scenario, the user does not need to supply any additional information to YAEL because anatomical information is automatically prepopulated in the electrode table.

**Figure 4.**
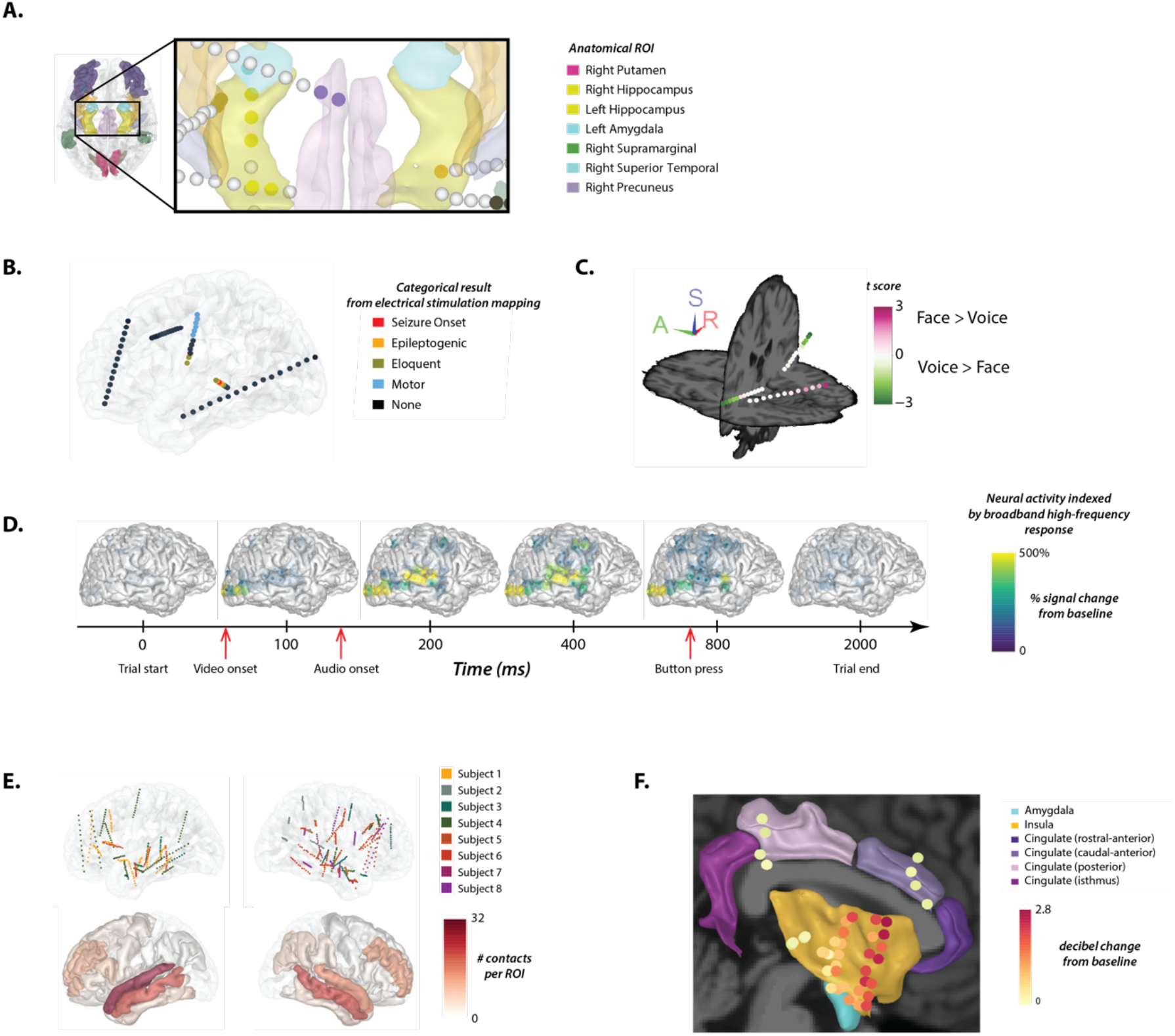
YAEL creates high-quality visualizations from a variety of iEEG data. A. YAEL visualization of anatomical data. Colored volumes show different anatomical regions-of-interest (ROI) from an atlas (legend at right). Spheres show sEEG electrode contacts. Colored contacts are located within an ROI, gray contacts are not in any ROI. B. YAEL visualization of categorical data. Contacts are colored by the results of electrical stimulation mapping. C. YAEL visualization of continuous data. Contacts are colored by the results of an analysis of iEEG power in two conditions, viewing faces and listening to voices. Only electrodes with a significant response are shown (non-significant contacts in gray). D. YAEL visualization of timeseries data. Each brain shows the power at one time point. Still frames show a movie where activity in each electrode was projected to the cortical surface. Alternately, only electrode activity can be shown, without projection to the cortical surface. Both movies available [here](https://openwetware.org/wiki/YAEL:Help#YAEL_Videos). E. YAEL visualization of group data. Top row shows electrodes from multiple participants, visualized on a template brain (one color per participant). Bottom row shows summary data of number of contacts across all participants in each anatomical ROI. F. YAEL visualization of combination data. Six anatomical regions were selected; all electrodes within each region across participants were selected; and then each electrode was colored by the iEEG response to a stimulus.

In the second scenario, YAEL is used to visualize categorical data about each electrode (Figure 4B). For instance, in electrical stimulation mapping current is applied through the implanted electrodes, one at a time, and the effect documented in a categorical way (George et al., 2020). An electrode might receive a designation of “motor” (if a motor response was evoked by stimulation), “eloquent cortex” if language function was interrupted, or “epileptogenic zone” if epileptiform activity was triggered. To create the data, the template table created by YAEL can be edited (using Microsoft Excel or another editor) to insert the appropriate values for each electrode; YAEL ignores missing values so it is not necessary to supply complete data; for instance, come electrodes could be omitted from stimulation mapping and the “stimulation results” column in the table would be left blank for these electrode rows.

In the third scenario, YAEL is used to visualize continuous data about each electrode, such as a clinical or research measure extracted from the iEEG data (Figure 4C). With continuous measures such as power, a common display mode is to select a statistically-significant threshold where only values above the threshold are displayed; if above threshold, the electrode is colored according to the continuous value. YAEL allows the user to independently choose the threshold and the color scale to use. For instance, the power in a given frequency band (time-locked to performance of a task or presentation of a sensory stimulus) provides information about the functional specialization of brain areas. Power may either decrease or increase relative to baseline, so power in response to a sensory stimulus (audiovisual speech) can be colored with a cold-to-hot color scale in YAEL. To create the data, the program used to calculate the continuous measure saves the data to a table, which YAEL then reads.

A fourth scenario is the display of time-series data (Figure 4D). When provided with time-series data, YAEL can use them to create movies of brain activity over time (sample movies available [here](https://openwetware.org/wiki/YAEL:Help#YAEL_Videos)). For instance, while viewing a talking face, visual areas might respond first when the face become visible, followed by auditory areas at the onset of the talker’s voice (Karas et al., 2019; Metzger et al., 2020). Instead of a single value per electrode, the analysis program saves multiple columns of data to the electrode table with one column per time-point, which is then read by YAEL to generate the movie.

A fifth scenario is visualizing data across multiple participants (Figure 4E). If electrodes from multiple individual subjects have been localized, YAEL can align the brains to the same template and display all electrodes together on the template brain. This permits users to understand the anatomical distribution of electrodes across participants and the sample size available in each ROI.

It is also possible to combine the different scenarios (Figure 4F). For instance, across all subjects, electrodes in selected anatomical regions could be selected for display. Then, the response to an experimental task in those electrodes could be mapped to a continuous color scale, providing a concise, single figure that summarizes a large quantity of group data (Metzger et al., 2023). This also enables new discovery. For instance, the data plotted in Figure 4F reveals an anterior-to-posterior gradient within the insular cortex, where anterior electrodes respond more strongly than posterior electrodes. In traditional workflows, this might be missed: in many workflows, all electrodes within a single ROI are collapsed, and the iEEG data from different ROIs are shown in a summary plot (Sakon and Kahana, 2022). This is because traditional workflows often lack the ability provided by YAEL to easily visualize the power in each electrode separately.

## Summary

Electrode localization is a critical step in iEEG research, but existing methods suffer from significant limitations, spurring the creation of YAEL. The most significant advantage of YAEL is the ability to create high-quality visualizations of the most common iEEG data, including anatomical parcellation, categorical data, continuous data, time series data, group data across participants, and combinations of these. Once visualizations are generated in YAEL, they can be exported in high-quality PDF format, editable in Adobe Illustrator, for use in presentations, manuscripts and grant proposals. Visualization is very important for increasing discovery, reproducibility and reliability (Unwin, 2020) and the availability of tools such as YAEL should accelerate the pace of iEEG research.

## Figures and Figure Legends

**Supplementary Figure 1 Legend.**
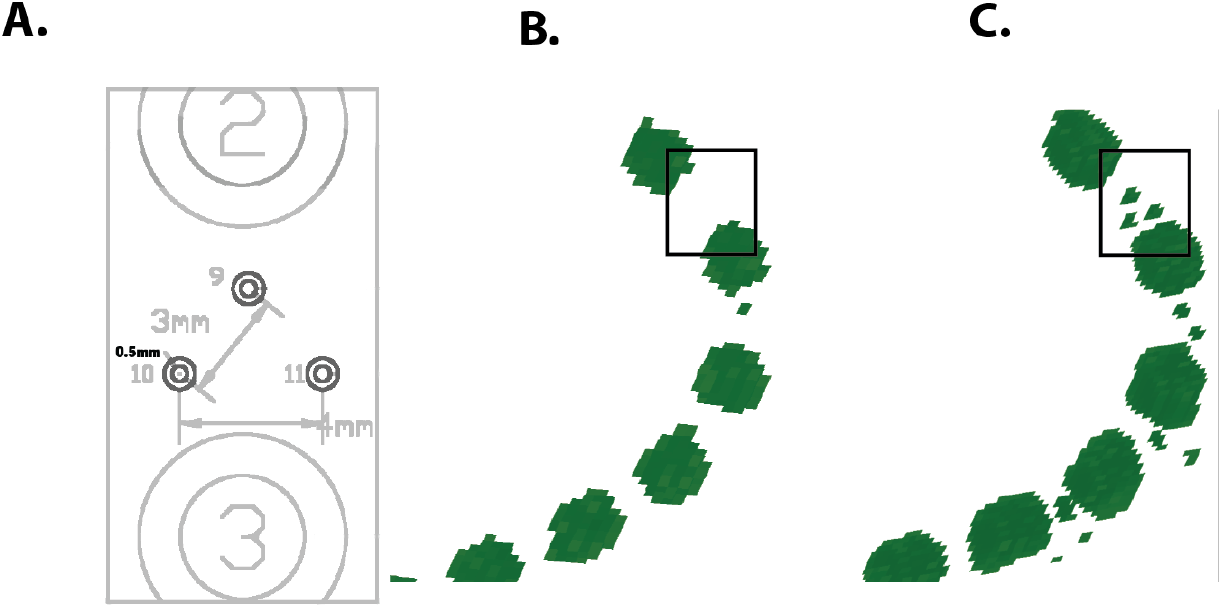
A. The manufacturer’s blueprint for an experimental subdural electrode strip, showing three miniature research contacts organized in a triangular pattern, located between two standard clinical contacts labelled “2” and “3”. B. If the CT is down sampled to the MRI resolution before viewing, the three miniature research contacts are not visible (black rectangle). C. In YAEL, the 3D viewer maintains the CT at the native resolution. The miniature research contacts are clearly visible (black rectangle).

## Acknowledgments

This research was supported by NIH R01NS065395 and U01NS113339.

## Notes

https://yael.wiki

### Competing Interest Statement

The authors have declared no competing interest.

